# Genomic evolution of SARS-CoV-2 variants of concern under *in vitro* neutralising selection pressure following two doses of the Pfizer-BioNTech BNT162b2 COVID-19 vaccine

**DOI:** 10.1101/2023.09.24.558921

**Authors:** Kerri Basile, Jessica E. Agius, Winkie Fong, Kenneth McPhie, Michael Fennel, Danny Ko, Linda Heuston, Connie Lam, Alicia Arnott, Sharon C-A Chen, Susan Maddocks, Matthew V. N. O’Sullivan, Dominic E. Dwyer, Vitali Sintchenko, Jen Kok, Rebecca J. Rockett, the CIDMLS COVID-19 Study Group

**Affiliations:** Centre for Infectious Diseases and Microbiology Laboratory Services, NSW Health Pathology - Institute of Clinical Pathology and Medical Research, Westmead Hospital, Westmead, New South Wales, 2145, Australia; Centre for Infectious Diseases and Microbiology – Public Health, Westmead Hospital, Westmead, New South Wales, 2145, Australia; The Westmead Institute for Medical Research, Westmead, New South Wales, 2145, Australia; Menzies Health Institute Queensland, Griffith University, Queensland, 4222, Australia; Sydney Institute for Infectious Diseases, Sydney Medical School, The University of Sydney, Westmead, New South Wales, 2145, Australia

**Author notes:** **Corresponding authors Dr Rebecca J. Rockett**, **Dr Kerri Basile**.

**Keywords:** Neutralising antibodies, SARS-CoV-2, VOC, B.1.167.2, B.1.351, COVID-19

## Abstract

**Aims:** To explore viral evolution during *in vitro* neutralisation using next generation sequencing, and to determine whether sera from individuals immunised with two doses of the Pfizer-BioNTech vaccine (BNT162b2) are as effective at neutralising the SARS-CoV-2 variant of concern (VOC) Delta (B 1.617.2) compared to the earlier lineages Beta (B.1.351) and wild-type (A.2.2) virus.

**Methods:** Using a live-virus SARS-CoV-2 neutralisation assay in Vero E6 cells we determined neutralising antibody titres (nAbT) in 14 participants (vaccine-naïve (n=2) and post-second dose of BNT162b2 vaccination (n=12), median age 45 years [IQR 29–65], median time after second dose = 21 days [IQR 19–28] against three SARS-CoV-2 strains: wild-type, Beta and Delta. The determination of nAbT was performed by visual inspection of cytopathic effect (CPE) and in-house quantitative reverse transcriptase real time quantitative polymerase chain reaction (RT-qPCR) to confirm SARS-CoV-2 replication. A total of 110 representative samples including inoculum, neutralisation breakpoints at 72 hrs, negative and positive controls underwent genome sequencing using the Respiratory Viral Oligo Panel version 2 (RVOP) (Illumina Inc. (San Diego, United States of America)) viral enrichment and short read sequencing using (Illumina Inc.San Diego, United States of America)(Figure 1).

**Results:** There was a significant reduction in nAbT observed against the Delta and Beta VOC compared with wild-type, 4.4-fold (*p* = >0.0006) and 2.3-fold (*p* = 0.0140), respectively (Figure 2). Neutralizing antibodies were not detected in one vaccinated immunosuppressed participant nor the vaccine-naïve participants (n=2). The highest nAbT against the SARS-CoV-2 variants investigated was obtained from a participant who was vaccinated following SARS-CoV-2 infection 12 months prior (Table S1). Limited consensus level mutations occurred in the SARS-CoV-2 genome of any lineage during *in vitro* neutralisation, however, consistent minority allele frequency variants (MFV) were detected in the SARS-CoV-2 polypeptide, spike (S) and membrane protein.

**Discussion:** Significant reductions in nAbT post-vaccination were identified, with Delta demonstrating a 4.4-fold reduction. The reduction in nAbT for the VOC Beta has been previously documented, however, limited data is available on vaccine evasion for the Delta VOC, the predominant strain currently circulating worldwide at the time. Studies in high incidence countries may not be applicable to low incidence settings such as Australia as nAbT may be significantly higher in vaccine recipients previously infected with SARS-CoV-2, as seen in our cohort. Monitoring viral evolution is critical to evaluate the impact of novel SARS-CoV-2 variants on vaccine effectiveness as mutational profiles in the sub-consensus genome could indicate increases in transmissibility, virulence or allow the development of antiviral resistance.

## Introduction

There remains ongoing worldwide spread of SARS-CoV-2, with variants of concern (VOC) arising independently in multiple locations worldwide. This, coupled with a notable increase in the substitution rate, suggests that positive selection of advantageous mutations is occurring within the SARS-CoV-2 genome. These mutations are particularly frequent in the spike (S) glycoprotein, the target for many vaccines and therapeutic antibody interventions [1,2]. Given this rapid vaccine development with roll out commencing 12 months following the first reported cases of COVID-19 with the Pfizer-BioNTech vaccine (BNT162b2), a nucleoside-modified RNA (mRNA) vaccine targeting the S protein in December 2020 [3,4]. This meant trials were largely conducted prior to the widespread circulation of VOC resulting in limited vaccine efficacy data against the VOC.

The constellation of mutations in the S glycoprotein of each VOC can reduce the effectiveness of natural and vaccine-induced protection [4–9]. The Delta (B.1.617.2) VOC possesses 12 mutations, most notably non-synonymous S mutations L452R, T478K, and P681R relative to the wildtype SARS-CoV-2, it lacks markers of convergent evolution such as mutations in S at amino acid positions N501Y or E484K/Q in its angiotensin-converting enzyme 2 (ACE2) receptor-binding domain [10]. The Delta VOC [11] does contain novel non-synonymous mutations within the S, such as T478K which has been previously described to decrease susceptibility to monoclonal antibody (mAb) treatment and acquired during persistent infection in an immunocompromised host [12].

Concerningly, this increased substitution rate coinciding with subsequent waves of infections and the roll out of COVID-19 vaccines globally, could be due to selective pressure from natural immunity, or have emerged through infection in immunosuppressed hosts [12,13]. Whilst genomic surveillance of SARS-CoV-2 can be used to monitor for new mutations, to predict the impact of these mutations on the efficacy of natural and or vaccine-derived immunity phenotypic assays are required.

Whilst live virus neutralisation remains the gold standard for determining antibody efficacy, [14] and neutralising antibodies (nAb) elicited by vaccination are considered correlates of protection from SARS-CoV-2 infection [15]. Reports of rapid evolution of SARS-CoV-2 during propagation in VeroE6 cells have emerged.

Including the generation of large genomic deletions removing the S1/S2 junction which encodes a putative furin cleavage site [29] and minority allele frequency variants (MFV) [50]. This cleavage site primes S for cell entry by exposing the S2 fusion peptide to enable virion fusion with the host cell membrane. Transmission of the virus with the deletion is attenuated in hamsters and ferrets but outgrows wild-type virus in VeroE6 cells [37,51]. This is of particular concern as VeroE6 cells are the predominate cell line used in studies reporting decreases in neutralising antibody titres (nAbT) of SARS-CoV-2 variants to sera from individuals post COVID-19 vaccination. Fold reductions in neutralisation reported from live virus micro-neutralisation assays performed in VeroE6 cells could therefore be overestimated by these common mutations occurring due to culture adaptation.

In this study we used sera collected from Australian health care workers after two doses of BNT162b2 to assess vaccine effectiveness when challenged during live virus infections with the Delta and Beta VOC compared with the wild-type strain. We sequenced the viral outgrowth to monitor the consensus and sub-consensus viral evolution *in vitro* to gain a greater understanding of genomic sites under evolutionary pressure or culture adaptation to potentially inform effective public health measures to limit the transmission of VOC.

## Methods

### SARS-CoV-2 culture

Upper respiratory tract specimens collected in universal transport media (UTM) where SARS-CoV-2 RNA was detected by real time reverse transcriptase real time polymerase chain reaction (RT-PCR) one either a Cobas^®^ 6800 (Roche Diagnostics GmbH (Mannheim, Germany)), a BD MAX™ (Becton Dickinson (Franklin Lakes, United States of America)), or an in-house assay [16] were used to inoculate Vero C1008 (Vero 76, clone E6, Vero E6 (ECACC 85020206), or Vero E6 expressing transmembrane serine protease 2 (TMPRSS2) cells [JCRB1819] cells as previously outlined [17].

In brief, cells were seeded at 1-3 × 10^4^ cells/cm^2^ whilst in the log phase of replication with Dulbecco’s minimal essential medium (DMEM) (BE12-604F, Lonza Group AG (Basel, Switzerland)) supplemented with 9% foetal bovine serum (FBS) (10099, Gibco™, Thermo Fisher Scientific Inc. (Waltham, United States of America)) in Costar^®^ 25 cm^2^ cell culture flasks (430639, Corning Inc. (Corning, United States of America)). The media was changed within 12 hrs for inoculation media containing 1% FBS and 1% antimicrobials (including amphotericin B deoxycholate (25 μg/mL), penicillin (10,000 U/mL), and streptomycin (10,000 μg/mL)) (17-745E, Lonza Group AG (Basel, Switzerland)) to prevent microbial overgrowth and then inoculated with 500 μL of clinical specimen into Costar^®^ 25cm^2^ cell culture flasks. Following inoculation of the clinical sample, all manipulation of SARS-CoV-2 cultures was performed under biosafety level 3 (BSL3) conditions [18].

Cultures were inspected daily for cytopathic effect (CPE); the inoculum and supernatant were sampled at 96 hrs for SARS-CoV-2 in-house quantitative reverse transcriptase real time polymerase chain reaction (RT-qPCR) targeting the *N*-gene as previously described [19]. A ≥ 3 cycle decrease in the cycle threshold (Ct) from the inoculum RT-qPCR result (equivalent to a one log increase in viral load - data not shown) as well as the presence of CPE was used to determine the propagation of SARS-CoV-2. Viral culture supernatant was harvested 96 hrs post-infection and a 500 μL aliquot was used to make a SARS-CoV-2 culture bank, where a large volume of passage one stock was made and stored at -80°C in 500 μL aliquots in 2 ml cryovials (72.694.406, Sarstedt Inc. (Nümbrecht, Germany)) until required, detailed in (Table 2). SARS-CoV-2 complete genomes were sequenced from the initial clinical specimen, positive culture supernatant, and passage one virus stock to quantify genomic variations that may have developed during propagation (Table S2).

### Human sera bank

Human sera was sourced from Australian healthcare workers caring for, or handling specimens from, individuals exposed to, or diagnosed with, SARS-CoV-2 infection enrolled in the COVID Heroes Serosurvey (http://www.covidheroes.org.au). Sera were tested upon receipt with an in-house immunofluorescence assay (IFA) against SARS-CoV-2 specific IgA, IgM and IgG [20] and then stored at -80°C. Fourteen sera samples were included from an age and sex matched cohort of 12 participants (Table S1). This included two vaccine-naïve individuals and 12 individuals who received two doses of BNT162b2 according to the schedule. Median age was 46 years [IQR 29–65] and median time after second dose of vaccine was 21 days [IQR 19–28]. Eleven of the 12 vaccine recipients had no documented history of prior SARS-CoV-2 infection, as confirmed by absence of SARS-CoV-2-specific antibodies on serial sampling since study enrolment. The remaining vaccine participant had laboratory confirmed SARS-CoV-2 infection one year prior to vaccination. Sera was heat-inactivated at 56 °C for 30 min to inactivate complement prior to microneutralisation.

### Determination of 50% tissue culture infective dose (TCID_50_)

The viral 50% tissue culture infective dose (TCID_50_) was determined for each variant virus. Briefly, a passage one aliquot of virus stock was serially diluted (1 × 10^−2^ – 1 × 10^−7^) in virus inoculation media. Virus dilutions were used to inoculate Vero C1008 (Vero 76, clone E6, Vero E6 [ECACC 85020206]) cells at 80% confluence in Costar® 24-well clear tissue culture-treated multiple well plates (Corning Inc. (Corning, United States of America)). Dilutions were seeded in duplicate with two negative (no virus) controls per plate. Plates were sealed with AeraSeal® Film (BS-25, Excel Scientific Inc. (Victorville, United States of America)) to minimise evaporation, spillage, and well-to-well cross-contamination. Plates were inspected daily for CPE and 100 μL sampled from each duplicate after inoculation and at 72 hrs. Infections were terminated at 72 hrs based on visual inspection for CPE and used in conjunction with RT-qPCR results to determine the TCID_50_ of each isolate.

### Micro-neutralisation assay

Vero C1008 (Vero 76, clone E6, Vero E6 [ECACC 85020206]) cells were seeded with DMEM (BE12-604F, Lonza Group AG (Basel, Switzerland)) from stocks in Costar□ 96-well clear tissue culture-treated flat bottom plates (353072) (Corning Inc. (Corning, United States of America)) at 40% confluence. Cells were incubated at 37 °C with 5% C0_2_ for 12 hrs or until they reached 80% confluence. Virus stocks were diluted to 200 TCID_50_ in inoculation media. Doubling dilutions from 1:10 to 1:320 of vaccine-naïve and post BNT162b2 vaccination sera were added in equal proportions with virus in a 96 well plate and incubated for 60 min at 37°C 5% C0_2_ to enable virus neutralisation. After this incubation the media was removed from the cell monolayer and 100 μL of fresh media was added. Each dilution of sera was performed in duplicate per virus variant, 12 wells of uninfected cells were used on each plate as a negative control. Plates were sealed with AeraSeal® Film to minimise evaporation, spillage, and well-to-well cross-contamination. After 60 mins of viral neutralisation a residual 110 μL was sampled from the 12 naïve patient wells per virus for extraction and RT-qPCR. The plates were inspected daily for CPE with a final read recorded at 72 hrs independently by two scientists. SARS-CoV-2 in-house RT-qPCR was used to quantify the viral load post-neutralisation, with 110 μL of each dilution removed at 72 hrs to determine viral load. The 110 μL of each dilution was added to 110 μL of External Lysis buffer (06374913001, Roche Diagnostics GmbH, (Mannheim, Germany)) at a 1:1 ratio in a 96-well deep-well extraction plate (Roche Diagnostics GmbH), covered with a MagNA Pure Sealing Foil (06241603001, Roche Diagnostics GmbH (Mannheim, Germany)), and left to rest in the biosafety class two cabinet for 10 mins, a time-period shown to inactivate SARS-CoV-2 by in-house verification of a published protocol [21]. The RNA was then extracted with the Viral NA Small volume kit (06 543 588 001, Roche Diagnostics GmbH (Mannheim, Germany)) on the MagNA Pure 96 system (Roche Diagnostics GmbH (Manheim, Germany)).

### SARS-CoV-2 genome sequencing following Respiratory Viral Oligo Panel enrichment

A total of 110 samples underwent SARS-CoV-2 whole genome sequencing. These included RNA extracts collected 1 hr post neutralisation (replicates of naïve 1:10 sera neutralisation) representing the baseline viral inoculum and neutralisation breakpoints as defined by CPE for each sera tested 72 hrs post neutralisation. Both biological replicates of each breakpoint were included, as were replicates of the naïve neutralisation at the highest sera dilution (1:10 and 1:20). Five specimens collected 72 hrs after neutralisation from uninfected wells on each plate were used as negative controls. A synthetic RNA SARS-CoV-2 construct (TWIST Biosciences) containing the reference SARS-CoV-2 sequence (National Center for Biotechnology Information (NCBI) GenBank accession MN908947.3) was diluted in negative control RNA (1:10) and was included in triplicate to control for library preparation and sequencing artefacts.

Viral enrichment was performed using the Illumina RNA Prep with the Respiratory Viral Oligo Panel version 2 (RVOP) (Illumina Inc. (San Diego, United States of America)) (Figure 1). RNA extracts from the microneutralisation and TCID_50_ experiments were used as input into the RNA Prep with Enrichment kit (Illumina Inc. (San Diego, United States of America)). RNA denaturation, first and second strand cDNA synthesis, cDNA tagmentation, library MFV construction, clean up and normalisation were performed according to manufacturer’s instructions. Individual libraries were then combined in 3-plex reactions for probe hybridisation. The RVOP was used for probe hybridisation with the final hybridisation step held at 58°C overnight. Hybridised probes were then captured and washed according to manufacturer’s instructions and amplified as follows: initial denaturation 98°C for 30 s, 14 cycles of: 98°C for 10 s, 60°C for 30 s, 72°C for 30 s, and a final 72°C for 5 mins. Library quantities and fragment size were determined using Qubit™ 1x dsDNA HS Assay (Invitrogen – ThermoFisher Scientific Inc. (Waltham, United States of America)) and Agilent HS D1000 Screentapes (Agilent Technologies Inc. (Santa Clara, United States of America)) Resulting libraries were pooled with the aim of generating 1×10^6^ raw reads per specimen and sequenced producing paired 74 base pair reads on the Illumina MiniSeq or iSeq instruments (Illumina Inc. (San Diego, United States of America)) (Figure 1).

**Figure 1.**
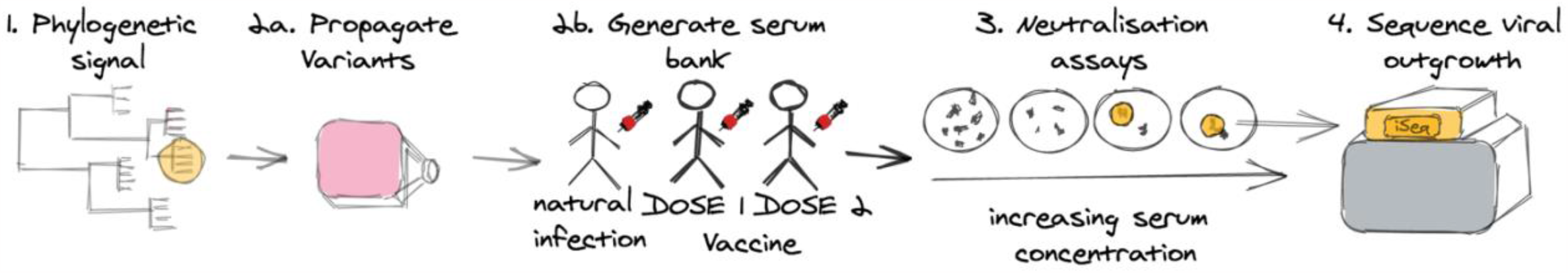
Outline of virus neutralisation assays of emerging SARS-CoV-2 variants of concern.

### Bioinformatic analysis

Raw sequence data were processed using an in-house quality control procedure prior to further analysis. De-multiplexed reads were quality trimmed using Trimmomatic v0.36 (sliding window of 4, minimum read quality score of 20, leading/trailing quality of 5 and minimum length of 36 after trimming) [22]. Briefly, reads were mapped to the reference SARS-CoV-2 genome (NCBI GenBank accession MN908947.3) using Burrows-Wheeler Aligner (BWA)-mem version 0.7.17 [23], with unmapped reads discarded. Average genome coverage was estimated by determining the number of missing bases (N’s) in each sequenced genome. Variants were called using VarScan v 2.3.9 [24] (min. read depth >10x, quality >20, min. frequency threshold of 0.1). Single nucleotide polymorphisms (SNP)s were defined based on an alternative frequency ≥0.75 whereas MFV were defined by an alternative frequency between 0.1 and 0.75. Variants falling in the 5’ and 3’ UTR regions were excluded due to poor sequencing quality of these regions. Polymorphic sites that have previously been highlighted as problematic were monitored and annotated in the results [25]. To ensure the accuracy of variant calls, only high-quality genomes with greater than 99% genome coverage and a median depth of 200x were included. The MFV calls were excluded in the base pair either side of the 5’ or 3’-end of indels due to miss-mapping. SARS-CoV-2 lineages were inferred using Phylogenetic Assignment of Named Global Outbreak LINeages v1.36.8 (PANGO).[26] Graphs were generated using RStudio (version 3.6.1).

### Statistical analysis

Mean nAbT were evaluated and statistical significance assessed using the t test with a 2 tailed hypothesis. Results were considered statistically significant at p < 0.05.

## Results

### Levels of neutralising antibody against different SARS-CoV-2 lineages

Genomic sequencing results indicated that the samples sequenced were wild-type strain (lineage A.2.2), Beta (lineage B.1.351) or Delta (lineage B.1.617.2) VOC. Following-vaccination with two doses of BNT162b2, 11 of 12 recipients demonstrated a functional neutralisation response to wild-type virus (Table S1 and Figure 1). The sera from an immunosuppressed participant that failed to mount a serological response to wild-type virus post-vaccination was excluded from further analysis to calculate fold reductions in nAbT (Table S1). No detectable antibodies or functional neutralisation responses were seen in sera collected prior to vaccination (n=2) (Table S1).

The median nAbT in BNT162b2 vaccine recipients who responded (n=11) when challenged with wild-type virus was 160 (range <10 – 320), compared to 80 (range <10 – 320) and 40 (range <10 – 80) for Beta and Delta respectively (Figure 2). There was a significant fold reduction in nAbT observed between both the Delta (*M* = 4.4, *SD* = 2), *t* (11) =-4.9, *p* = 0.00059, and Beta (*M=*2.3, *SD=*2) *t* (11) =-3 *p* = 0.01397 compared with wild-type (Figure 2). There was also a significant fold reduction in nAbT between Beta (*M* = 2.6, *SD* = 1.4), *t* (11) =-2.5, *p* = 0.02897 and Delta (Figure 2).

**Figure 2:**
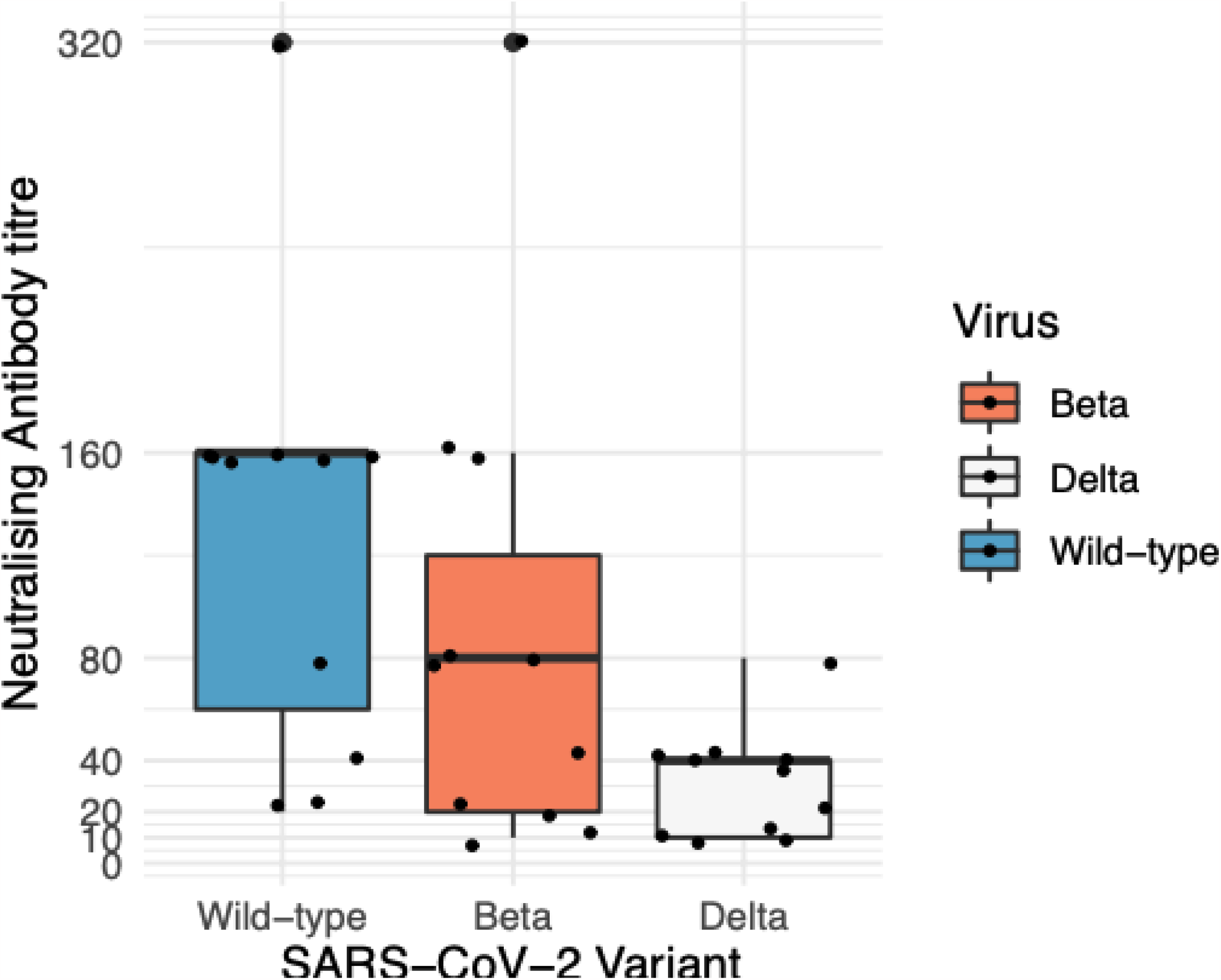
Neutralising Antibody Titres. **Figure description:** Differences in neutralising antibody titres (nAbT) against the Wildtype, Beta and Delta SARS-CoV-2 lineages median of 21 days (IQR 19-28) after receiving the second dose of Pfizer-BioNTech (BNT162b2) vaccine measured by visual inspection of cytopathic effect (CPE) with confirmation of viral replication by SARS-CoV-2 by inhouse reverse transcriptase real time quantitative polymerase chain reaction (RT-qPCR). Results are reported in the box−whiskers plots as medians and upper and lower quartiles. There was a significant fold reduction in nAbT observed between both the Delta (*M* = 4.4, *SD* = 2), *t*(11) =-4.9, *p* = .00059, and Beta (*M=*2.3, *SD=*2) *t*(11) =-3 *p* = .01397 compared with wild-type. There was also a significant fold reduction in nAbT between Beta (*M* = 2.6, *SD* = 1.4), *t*(11) =-2.5, *p* = .02897 and Delta. **Key:** Delta – Delta (B.1.617.2) lineage; Beta – Beta (B.1.351) lineage; Wildtype – Wildtype (A.2.2) lineage;

Participants aged ≤ 45 years had a significantly higher fold reductions in nAbT for (*M = 5*.*6, SD = 2*.*2*) compared with those aged > 46 (*M=3*.*3, SD=1*) between the wild-type and Delta strains *t* (5) =4.9 *p* =0 .00788. The ≤ 45 years cohort also had significantly higher fold reductions in nAbT *M =* 4, *SD =* 0) compared with >46 yrs (*M=*1.4, *SD*=0.7) between Beta and Delta *t* (5) =-3.2, *p* =0 .03397. No significant gender specific effects were identified. The highest nAbT for all SARS-CoV-2 variants investigated was obtained from a participant who was infected with SARS-CoV-2 one year prior to vaccination (Table S1).

### Quantification of inoculation dose for different lineages of SARS-CoV-2

The mutational profile of the inoculum (Table S2 and S3) for each lineage was consistent with the mutational profile defining the PANGO lineages assigned to the original clinical specimen (Figure 3). The SARS-CoV-2 genomes of the inoculated viruses, the original clinical specimens and viruses isolated in cell culture were compiled with representation of the global SARS-CoV-2 diversity (n=1000) curated by Nextstrain (Nextstrain.org) (Figure 3). The viral stock used had an infecting dose of 200 TCID_50_. The Ct value of the inoculum of the wildtype and Beta VOC was 28, whereas the Ct of the Delta was 24. This difference in Ct values could be explained by the increased sensitivity of PCR assays, however PCR is unable to differentiate between infectious SARS-CoV-2 virions and non-viable virus.

**Figure 3.**
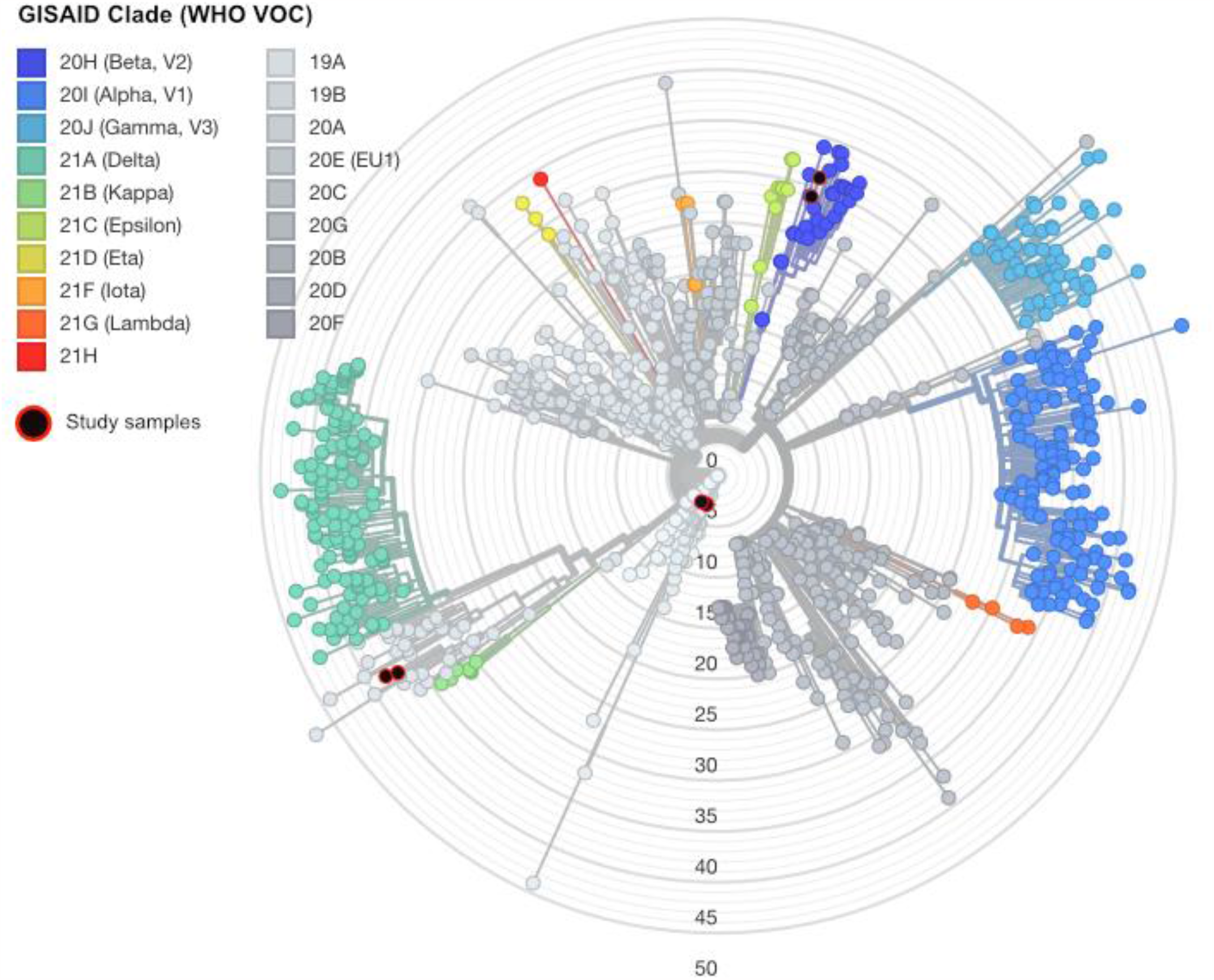
Global SARS-CoV-2 diversity demonstrating representativeness of inoculating viruses used in this study. **Figure Description:** The genome sequence of the SARS-CoV-2 lineages neutralised by sera post complete vaccination with BNT162b2 were included in the subsampled global phylogeny of SARS-CoV-2. Representative sequences were selected by Nextstrain and used to generate a global phylogeny, the original clinical specimen and inoculating virus sequence is highlighted in black with red outline. **Key:** VOC – variant of concern; WHO – World health organization;

### SARS-CoV-2 polymorphism in culture

A total of 110 samples underwent SARS-CoV-2 sequencing including 102 extracts from the microneutralisation experiment, five negative controls, and the three SARS-CoV-2 synthetic constructs as positive controls (TWIST Biosciences, encoding NCBI GenBank accession MN908947.3). All but one genome was recovered with high read depth (average depth 3300.5x (range 174 - 11844) and the average genome coverage was 99.97% (range 99.73 - 100%). The five negative control samples contained <10 SARS-CoV-2 specific reads.

A total of 3039 polymorphisms were detected during neutralisation (majority allele variants =2715 and MFV = 324), and the highest frequency base change was C>U (Figure S2). We focused our investigation on genomic variants that were not in the viral inoculum and developed 72 hrs post-neutralisation (majority allele variants = 21 and MFV = 176). Base change dynamics were similar between majority variant polymorphisms, *de novo* consensus level changes and *de novo* MFV, apart from G>C changes noted at high frequency in the *de novo* MFV (Figure S2). Non-synonymous mutations were detected at greater frequency than synonymous, indels and nonsense mutations. A higher ratio of non-synonymous to synonymous (Ka/Ks) mutations was detected when comparing *de novo* MFV (Ka/Ks 3.35) to majority frequency variants (Ka/Ks 1.38) (Figure S2).

Aliquots of the viral-sera inoculum were collected 1-hour post-neutralisation in biological replicates for each SARS-CoV-2 variant investigated. The consensus and MFV mutations were tabulated and used as a baseline for our analysis (Table S3).

No additional mutations or MFV were noted in the viral inoculum within the furin cleavage site, previously reported after long term passage in VeroE6 cells (Table S3)[27–33].

### Persistence and conversion of minority frequency allele variants 72hours post-neutralisation

Minimal consensus level mutations were noted 72 hrs post-neutralisation compared to the viral inoculum (Figure 4 and Supplementary Figure 3); however, sub-consensus MFV persisted in Beta and Delta infections (Supplementary Figure 3). Persistence of MFV at position C13667T (nsp12, orf1ab p.4468T>I) detected at a read frequency of 0.06 - 0.22 was noted in 32/33 biological replicates, including the inoculating virus. A total of six MFV were detected in the Beta inoculum, four of which persisted and generally increased in frequency 72 hrs post-neutralisation. The MFV at position C11249T (nps6, orf1ab p.3662R>C) was detected 1-hour post-neutralisation at an average frequency of 0.53, the mutation persisted in 26/32 replicate infections at 72 hrs and developed into a consensus base change in 13/26 replicates. The MFV at position C11750T (nsp6, orf1ab p.3829L>F) was detected 1-hour post-neutralisation at a median frequency of 0.25 and persisted in 20/32 infections at 72 hrs, developing into consensus mutations in 6 infections. Synonymous mutations at C27911T (orf8 p.6F) were detected at an average frequency of 0.065 1-hour post-neutralisation, and persisted in 9/32 infections, converting to a consensus mutation in a single infection. A second synonymous MFV at position C29077T (N: p.268Y) was detected at a frequency of 0.66 1-hour post-neutralisation and was detected in 27/32 infections at 72 hrs converting to a SNP in 12/27 infections. However, the two MFV detected 1-hour post-neutralisation for the wild-type variant (G18670T, nsp14, orf1ab p.6136D>Y, G22316A S: p.252G>S) were detected at low read frequency (<0.01) and did not persist in any infections at 72-hrs.

**Figure 4.**
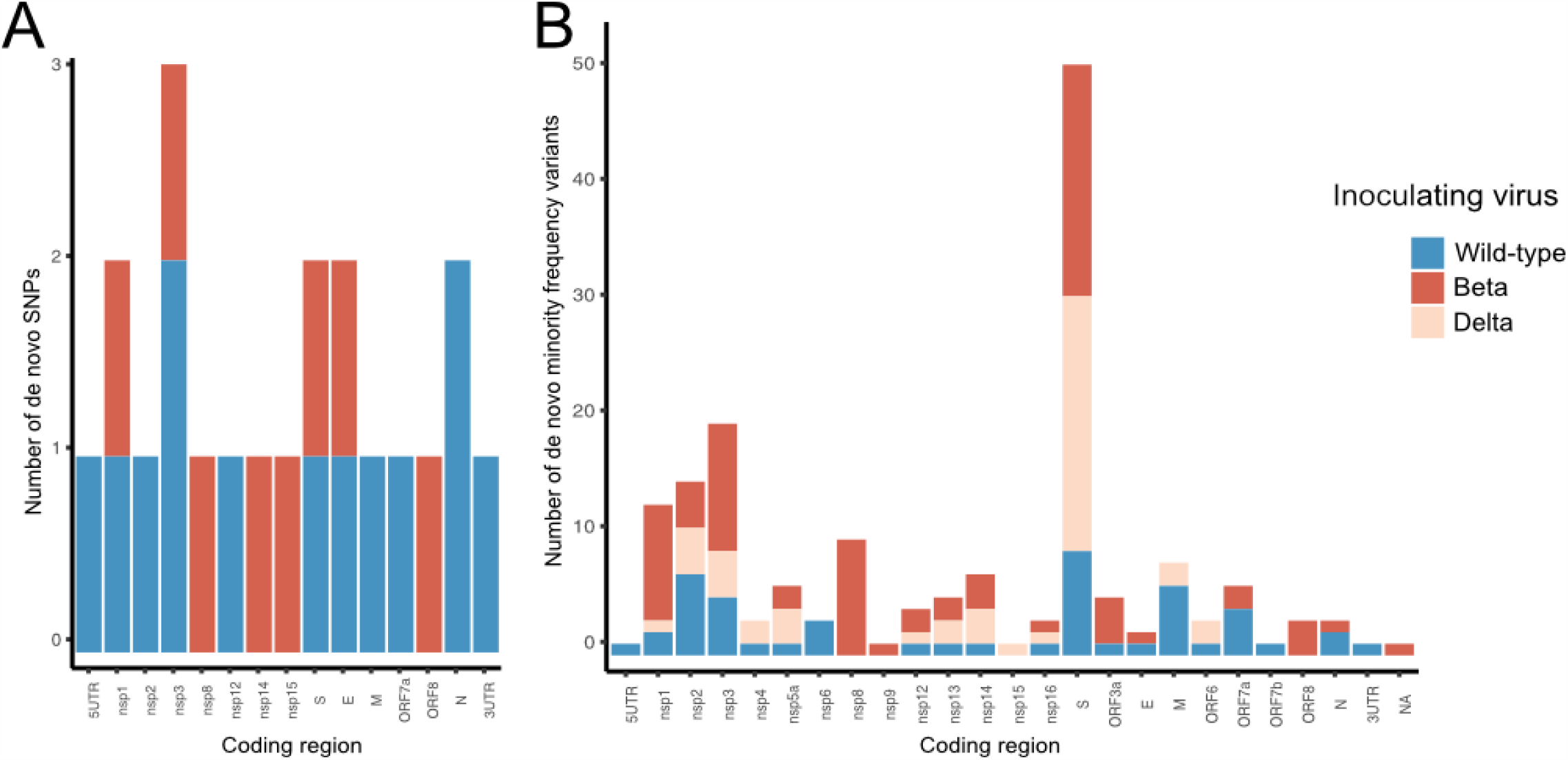
Frequency of consensus and minority allele frequency variants within the SARS-CoV-2 genome. **Figure Description:** Count of *de novo* consensus (**A**) and minority frequency allele variants (MFV) (**B**) within SARS-CoV-2 genes 72 hours post neutralisation. Consensus and MFV detected in the inoculating virus of each lineage have been excluded. Although few consensus level mutations were detected 72 hours post-neutralisation, a high number of MFV were detected within the spike (S) coding region. **Key:** structural proteins [S - spike ; E – envelope; M – membrane and N – nucleocapsid]; nsp – non-structural protein; ORF– open reading frame; NC – non-coding region, 5UTR – 5’ untranslated region, 3UTR – 3’ untranslated region, Delta – Delta (B.1.617.2) lineage; Beta – Beta (B.1.351) lineage; Wildtype – Wildtype (A.2.2) lineage; Dose 2 – post the 2^nd^ dose of Pfizer-BioNTech (BNT162b2) SARS-CoV-2 vaccination; Reference – reference SARS-CoV-2 genome (NCBI GenBank accession MN908947.3)

### De novo majority frequency variants

*De novo* mutations that were not detected 1-hour post-neutralisation were also investigated (n=21, Figure 4, Supplementary Figure 2). When wildtype virus (A.2.2) was neutralised, a maximum of three consensus level mutations developed in any infection compared to the sequence of the inoculum (median 0, range 0 - 3) (Figure 3 and 4, Table S3). Of 11 consensus level mutations within the coding region, six were synonymous and five were non-synonymous (Figure S2). Only eight *de novo* consensus level mutations were detected 72 hrs post-neutralisation after inoculating with the Beta variant. A maximum of two consensus level mutations developed in any infection compared to the inoculum sequence (median 0, range 0-1). None of the genomic positions detected were replicated over the 32 infections. No consensus level mutations developed 72 hrs post infection when Delta was neutralised in the 34 infections.

### Development of de novo minority allele frequency variants

MFV were generally homogeneously detected across the SARS-CoV-2 genome; however, a concentration of MFV in the S protein was noted (Figure 4). Novel MFV (n=63) were detected at 32 unique genomic positions 72 hrs post-neutralisation of the wild-type virus (Figure 4). Three of these genomic locations (C2156T nsp2 orf1ab p.631L>F, G25337C S:p.1258D>H, C26895T M:p.125H>Y) were reproducibly detected in >5 infections (Figure 5). The MFV converted to a consensus change at position C2156T in a single infection.

**Figure 5.**
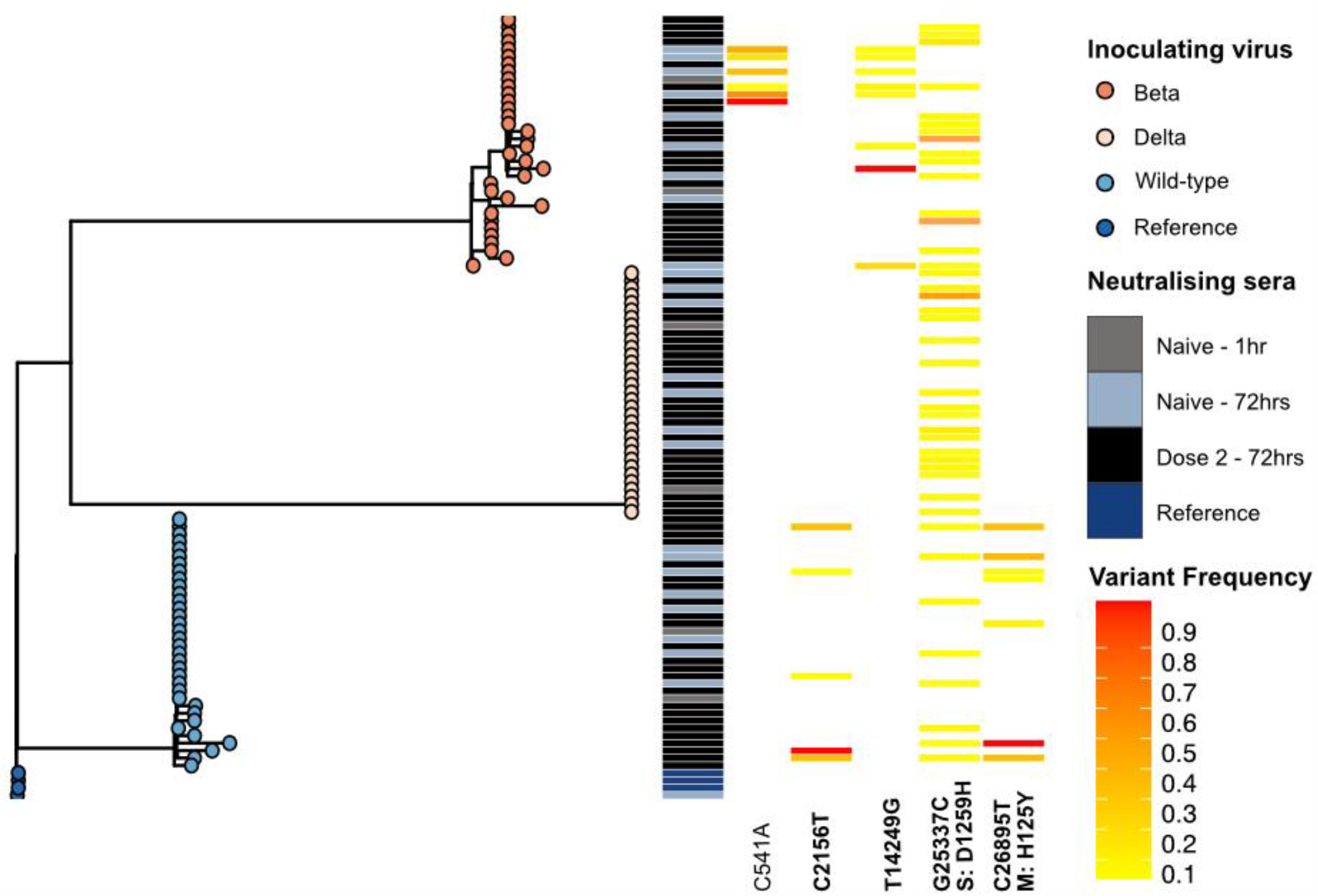
Phylogeny of SARS-CoV-2 diversity during neutralisation highlighting minority variants reproducibly detected at the same genomic location. **Figure Description:** A maximum likelihood phylogeny of the 109 SARS-CoV-2 genomes generated in this study. Tree node colours indicate the lineage of the inoculating virus and the heatmap highlights the time of sampling and the vaccine dose of the sera used. *De novo* minority allele frequency variants (MVF) which were consistently detected (≥ 5 biological replicates) at specific genomic locations are shown in the heatmap. The read frequency of these sub-consensus variants is depicted by the colour scale, where a frequency of 0.1 is shown in yellow and a frequency of 0.9 is shown in red. The MVF variants detected in the inoculating viruses were not included. **Key:** Delta – Delta (B.1.617.2) lineage; Beta – Beta (B.1.351) lineage; Wildtype – Wildtype (A.2.2) lineage; 72 hrs– samples collected at 72 post neutralisation, Clinical sample – from the original patient’s sample used to generate the culture isolate.

Novel MFV (n= 86) were detected in 50 unique position 72 hrs post-neutralisation of the Beta VOC. Three genomic positions ((C541A nsp1 orf1ab p.92L), (T14249G nsp8, orf1ab p.4662L>W), (G25337C S:p.1258D>H)) were reproducibly detected in >5 biological replicates. The MFV converted to a consensus change at position T14249G in a single infection.

When the Delta VOC was sequenced 72 hrs post-neutralisation, novel MFVs (n= 48) were detected at 28 unique genomic positions. A single MFV at position G25337C was repeatedly detected in 18/32 infections.

A high frequency of MVF variants in the S protein coding region were detected during genome-wide analysis. Of the 51 MVF variants detected in the S protein 72 hrs post-neutralisation with any SARS-CoV-2 virus, 41/51 were detected at position 25337 which encodes a non-synonymous mutation D1259H in the C-terminal domain of the S protein. This mutation was not detected in the genomes generated one-hour post-neutralisation. Only three MVF variants were detected within the S1/S2 cleavage site at nucleotide positions 23606 (S p.682R>W), 23616 (S p.691S>A), and 23633 (S p.697M>T) at read frequencies of 0.05, 0.09 and 0.08 respectively.

The only MFV that persisted across all lineages was at position G25337C. This variant was detected in 41/102 infections but was not detected in the three viral inocula. The MFV persisted at a low frequency (median 0.08 min 0.05 max 0.15)

Indels detected in the minority of reads were also uncovered in 9/102 infections, generating six deletions and four insertions. A 10bp deletion was detected at nucleotide position 685 (AAAGTCATTT685A, orf1ab p.141-144KSFD>X) 72 hrs after the Beta VOC was neutralised in two infections at a read frequency of 0.06 and 0.17. Two additional deletions were detected in nsp1 at positions 514 and 515 ((TGTTATG515T orf1ab p.84-85VM>X) and (GTTA515G orf1ab p.84-85VM>-)). Single base insertions and deletions were identified in single infections ((G1772GT orf1ab p.503V>F), (TG16911T orf1ab p.5550V>X), (T25878TC ORF3a p.162-163->X), (GA27396G ORF7a p.2K>X) (G27906GT ORF8 p.5V>VX).

## Discussion

The rapid and widespread global adoption of genomic sequencing combined with traditional epidemiology, to address the COVID-19 pandemic has enabled real-time surveillance of SARS-CoV-2 evolution. The ability to determine the frequency of cases with specific mutational profiles has provided strong indicators of SARS-CoV-2 variants with selective advantage [34–36]. Supporting evidence from *in vitro* studies, and SARS-CoV-2 coding positions demonstrating convergent evolution, has clearly highlighted mutations in the S protein that have increased infectivity [33,37].

In this study we confirm a reduction in nAbT after BNT162b2 vaccination when challenged with the Beta VOC [38–40], and demonstrated a significant 4.4-fold reduction in nAbT against the Delta VOC. Herein we highlight the higher fold reduction in nAbT against the Delta compared to the Beta VOC. Concordant with other studies, we demonstrate that sera from a vaccine recipient who was previously infected with SARS-CoV-2 mounted the highest nAbT against all virus variants [41] and immunocompromised individuals may fail to mount neutralising antibody responses after completing a two-dose BNT162b2 vaccination schedule [43]. Furthermore, assessments of vaccine effectiveness in populations with higher incidences of COVID-19 may not apply to populations with lower rates of COVID-19, such as Australia [42].

To ensure the validity of our findings and to control for genomic adaptations in the furin cleavage site, commonly reported when SARS-CoV-2 is cultured in VeroE6 cells, we undertook genomic analysis of the viral inoculum and outgrowth compared with the original clinical isolate.

At 72 hrs post-neutralisation limited *de novo* consensus mutations were noted when compared to the infecting VOC. However, several *de novo* MFV were detected in the inoculum and persisted in biological replicates post neutralisation demonstrating the utility of deep sequencing and hybridisation probe capture [44] to accurately monitor MFV. The higher resolution provided by sequencing enabled accurate monitoring of MFV but is confounded by homoplasy in the SARS-CoV-2 genome, transmission bottlenecks and the transient nature of many MFV during the course of infection [44–46].

With one exception these genomic locations were not conserved between infecting viral lineages, and the genomic positions were not in identified homoplastic sites. One MFV in the C-terminal of S was detected in 41/96 infections 72 hrs post-neutralisation. This MFV results in a D1258H mutation, which has only been reported in 50 SARS-CoV-2 consensus sequences available on the Global Initiative on Sharing All Influenza Data (GISAID) EpiCoVô database as of 12 November 2021[47]. Rocheleau *et al* reported the detection of this MFV in both clinical and cultured SARS-CoV-2 genomes and provides evidence that missense mutations that truncate the C-terminal domain of the S protein enable more efficient viral exocytosis by promoting direct cell-cell fusion. [48] The reproducible detection of this MFV requires further investigation to determine if it provides a selective advantage, or if this genomic site is homoplastic.

The importance of SARS-CoV-2 diversity driven by *de novo* mutations that occur within hosts is of great importance, particularly as evidence of positive selection of mutations that can evade immunity and therapeutics and have been demonstrated in immunocompromised patients and in *in vitro* systems [13,44,49].

The use of *in vitro* systems is critical for understanding population dynamics of SARS-CoV-2 infections as mutational profiles in the sub-consensus genome may potentially inform surveillance for variants that increase transmissibility, virulence and/or antiviral and mAb resistance, with controls in place for mutations driven by culture adaptation that may not correlate *in vivo*. Despite the relatively low number of participants studied herein, the cohort included age and sex matched participants being immunised with BNT162b2 exactly three weeks apart and a median date of sera collection post dose two of 21 (IQR21-29) to control for expected peak immunity post vaccination. The lack of detectable antibodies reflects the relatively low incidence of COVID-19 infection and initial slow vaccine uptake in the Australian population [42].

## Conclusion and future directions

More studies assessing convalescent samples and other markers of immunity to determine correlates and duration of protection against emerging variants of interest and VOC, as well as assessing nAbT responses in serially collected samples are required. Similar approaches could also be used for sera from individuals receiving other mRNA, viral vector, protein or inactivated SARS-CoV-2 vaccines [52]. Comparisons between primary and booster vaccinations should also be investigated to determine the optimal vaccination strategy at an individual (including those that are immunocompromised) and population level. In conclusion there is a significant reduction in nAb against the Delta compared to Beta and wild-type variants. Modelling has predicted that a five-fold decrease in nAbT would likely reduce the effectiveness of current vaccines from 95% to 77% for high efficacy vaccines and 70% to 32% for lower efficacy vaccines [53]. Deep sequencing and hybridisation probe capture, an approach less hampered by amplification biases, enables high resolution monitoring of SARS-CoV-2 evolution including mutational profiles in the sub-consensus genome. This is a critical factor in evaluating vaccine effectiveness as population level vaccination is underway. to evaluate as mutational profiles in the sub-consensus genome could indicate increases in transmissibility and or virulence.

## Acknowledgements

The authors would like to acknowledge the Sydney Informatics Hub, a Core Research Facility of the University of Sydney, for the use of high-performance computing infrastructure. We would also like to acknowledge the COVID Heroes (A serosurvey of healthcare workers caring for, or handling specimens from, individuals exposed to, or diagnosed with, SARS-CoV-2 infection in Australia) study participants from donation of sera samples. All laboratories who referred samples to the Centre for Infectious Diseases and Microbiology Laboratory Services, NSW Health Pathology - Institute of Clinical Pathology and Medical Research, Westmead Hospital, Westmead NSW 2145, Australia for testing that were included in this analysis.

## Ethics

Ethical and governance approval for the study was granted by the Western Sydney Local Health District Human Research Ethics Committee (2020/ETH02426) and (2020/ETH00786)

## Author contributions

|  |  |
| --- | --- |
| **Designed experiments** | KB, KM, DD, MO, VS, JK and RR |
| **Performed experiments** | KB, JA, WF, KM, LH, HR, MF, DK, TT, SK, IC, TS, LD, SA and RR |
| **Data analysis and visualisation** | KB, JA, WF, AA, KM CL, JD, MG, EM, SC, SM, DD, MO, VS, JK and RR |
| **Performed bioinformatic analyses** | KB, JA, WF, CL, JD, MG, EM and RR |
| **Logistics and resources** | KB, KM, LH, AA, HR, TT, LD, EM, CL, CN, JN, BS, RB, NJ, NR, JT, KK, HV, MO, VS, JK and RR |
| **Drafting the initial manuscript** | KB and RR |
| **Critical manuscript revision and supervision of research** | KB, SC, SM, DD, MO, VS, JK and RR |

