## Supplementary material for "Genomic evolution of SARS-CoV-2 variants of concern under *in vitro* neutralising selection pressure following two doses of the Pfizer-BioNTech BNT162b2 COVID-19 vaccine"

#### Table S1: Study participant demographic and inhouse SARS-CoV-2 immunofluorescence and neutralising antibody titres

| | Sex | Age (yrs) | Post 2nd dose BNT162b2 (days) | SARS-CoV-2 IFA | | | Lineage | | | Other |
|---|---|---|---|---|---|---|---|---|---|---|
| | | | | IgG | IgA | IgM | Wild-type | Beta | Delta | |
| **1** | F | 36 | 21 | 1280 | <10 | <10 | 160 | 80 | 20 | |
| **3** | M | 49 | 28 | 320 | <10 | <10 | 20 | <10 | <10 | |
| **4** | M | 29 | 22 | 640 | 10 | <10 | 80 | 40 | <10 | |
| **5** | M | 65 | 22 | 640 | <10 | <10 | 160 | 20 | 40 | |
| **6** | F | 35 | 21 | 1280 | 10 | <10 | 320 | 320 | 80 | Previously infected with SARS CoV-2 |
| **7** | F | 44 | 19 | <10 | <10 | <10 | <10 | <10 | <10 | Immunosuppressed |
| **8** | F | 45 | 21 | 640 | <10 | <10 | 40 | 20 | <10 | |
| **9** | F | 59 | 21 | 160 | <10 | <10 | 20 | <10 | <10 | |
| **10** | F | 36 | 21 | 640 | <10 | 10 | 160 | 160 | 40 | |
| **11** | F | 57 | 21 | 1280 | <10 | <10 | 160 | 80 | 40 | |
| **12** | M | 43 | 28 | 1280 | <10 | 10 | 160 | 160 | 40 | |
| **13** | M | 47 | 22 | 1280 | <10 | <10 | 160 | 80 | 80 | |
| **4** | M | 29 | naïve | <10 | <10 | <10 | <10 | <10 | <10 | Naïve pre vaccination sera |
| **11** | F | 57 | naïve | <10 | <10 | <10 | <10 | <10 | <10 | Naïve pre vaccination sera |

**Key:** Beta – Beta (B.1.351) lineage; BNT162b2 – Pfizer-BioNTech (BNT162b2) vaccination; Delta – Delta (B.1.617.2) lineage; F – female; IFA – Immunofluorescence Assay; IgA – immunoglobulin A; IgG – immunoglobulin G; IgM – immunoglobulin M; M – male; Wild-type – Wildtype (A.2.2) lineage;

### Table S2: Summary of viral stock used

| Lineage | WHO Variant of Concern | GISAID ID of original clinical isolate | Cells originally isolated in | Passage number |
|---|---|---|---|---|
| A.2.2 | - | hCoV-19/Australia/NSW183/2020 | Vero C1008 (Vero 76, clone E6, Vero E6 [ECACC 85020206] | 1 |
| B.1.351 | Beta | | TMPRSS2 expressing VeroE6 [JCRB1819] | 1 |
| B.1.617.2 | Delta | hCoV-19/Australia/NSW1601/2021 | TMPRSS2 expressing VeroE6 [JCRB1819] | 1 |

**Key:** GISAID – Global initiative on sharing all influenza data; Lineage – PANGO Lineage_ version 3;TMPRSS2 – transmembrane serine protease 2

#### Table S3: Mutational profile of inoculum

| Inoculating virus | Consensus mutations<br><br>(Protein changes) | Low frequency variants (read frequency, protein changes) |
|---|---|---|
| **Wild type (Lineage A.2.2)** | C4540T, C8782T, T9477A (nsp4, F308Y), C14805T, G25979T (ORF3a, G196V), T28144C (ORF8, L84S), C28311T (N, P13F), C28657T, C28863T (S197L) | G18670T (6.08%*, nsp14, D211Y), G22316A (5.37%*, S, G252S) |
| **Beta (Lineage B.1.351)** | G174T, C241T, C1059T (nsp2, T85I), C3037T, T4213C, G5230T(nsp3, K837N), A10323G (nsp5, K90R), C11812A, GTCTGGTTTT11287G, C14408T (nsp12, P323L), C15925T (nsp12, L829F), T17970C, A20662G (nsp16, S2G), G21641T (S, A27S), A21801C (S, D80A), A22206G (S, D215G), ACTTTACTTG22280A, G22813T (S, K417N), G23012A (S, E848K), A23063T (S, N501Y), A23403G (S, D614G), C23664T (S, A701V), A23764T, G25563T (ORF3a, Q57H), C25904T(ORF3a, S171L), C26456T (E, P71L), C28253T, T28642C, G28871T (N, G200C), C28887T (N, T205I) | C11249T (49%, nsp6, R97C), C11750T (25%, nsp6, L206F), C12114T (12%*, nsp8, S8F), T14637A (11%*), A27452G (5%*, ORF7a, Y20C), C27911T (7%),<br><br>C29077T (67%) |
| **Delta (Lineage B.1.617.2)** | G210T, C241T, C1191T (nsp2, P129L), C1267T, C3037T, G4201T (nsp3, M494L), C5184T(nsp3, P822L), C6539T (nsp3, H1274Y), C9891T (nsp4, A446V), T11418C (nsp6, V149A), T12946C, C14408T (nsp12, P323L), T15225C, G15451A (nsp12, G671S), C16466T (nsp13, P77L), A20262G, C20320T (nsp15, H234Y), C21618G (S, T19R), G21793T (S, K77B), G21987A (S, G142D), GAGTTCA22028G, C22591T, T22917G (S, L452R), C22995A (S, T478K), A23403G (S, D614G), C23604G (S, P681R), G24410A (S, D950N), C24745T, C25469T (ORF3a, S26L), C26353T (E, L37F), T26767C (M, I82T), T27638C (ORF7a, V82A), C27739T (ORF7a, L116F), C27752T (ORF7a, T120I),<br><br>AGATTTC28247A, TA28270T,<br><br>A28461G (N, D63G), G28881T (N, R203M), G29402T (N, D377Y), G29427A (N, R385K), G29742T | C13667T (14.18%, nsp12, T76I) |

#### Figure S1: Change in SARS-CoV-2 viral load three days post neutralisation in responders and non-responders

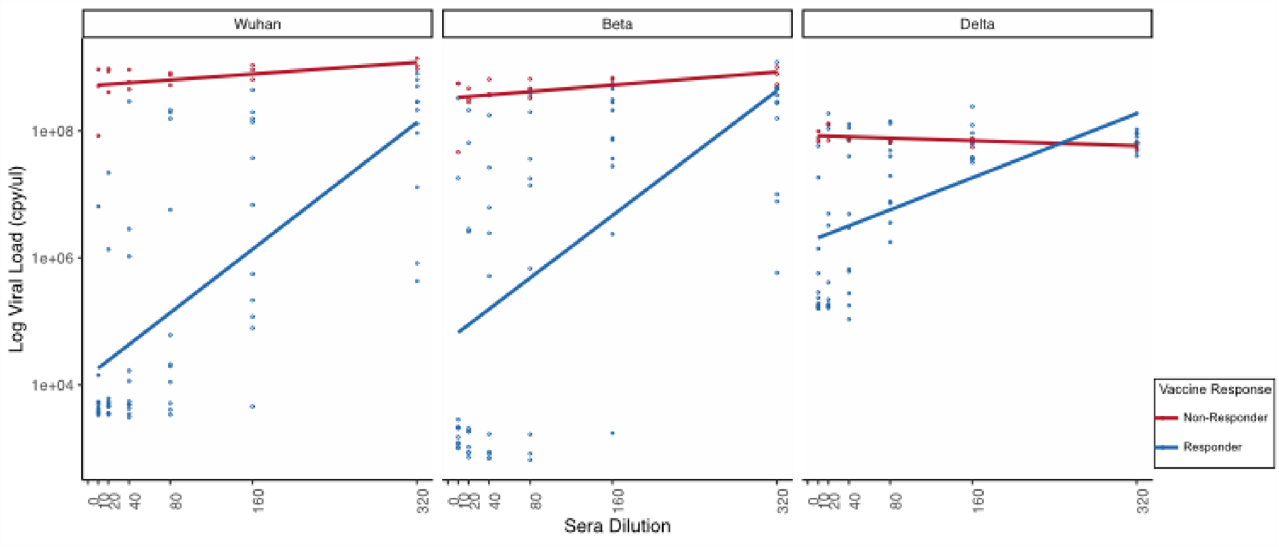


**Key:** Beta – (B.1.351) lineage; Delta - (B.1.617.2) lineage; Non- Responder (n=3) including naïve (n=2) and an immunocompromised patient post 2nd dose of Pfizer-BioNTech (BNT162b2) SARS-CoV-2 vaccination (n=1). Responders (n= 11) participants who had neutralising antibody titres detected post the 2nd dose of Pfizer-BioNTech (BNT162b2) SARS-CoV-2 vaccination; Wild type – (A.2.2) lineage;

**Figure S2: Mutational analysis of fixed mutations and minority allele frequency variants during live virus *in vitro* neutralisation**

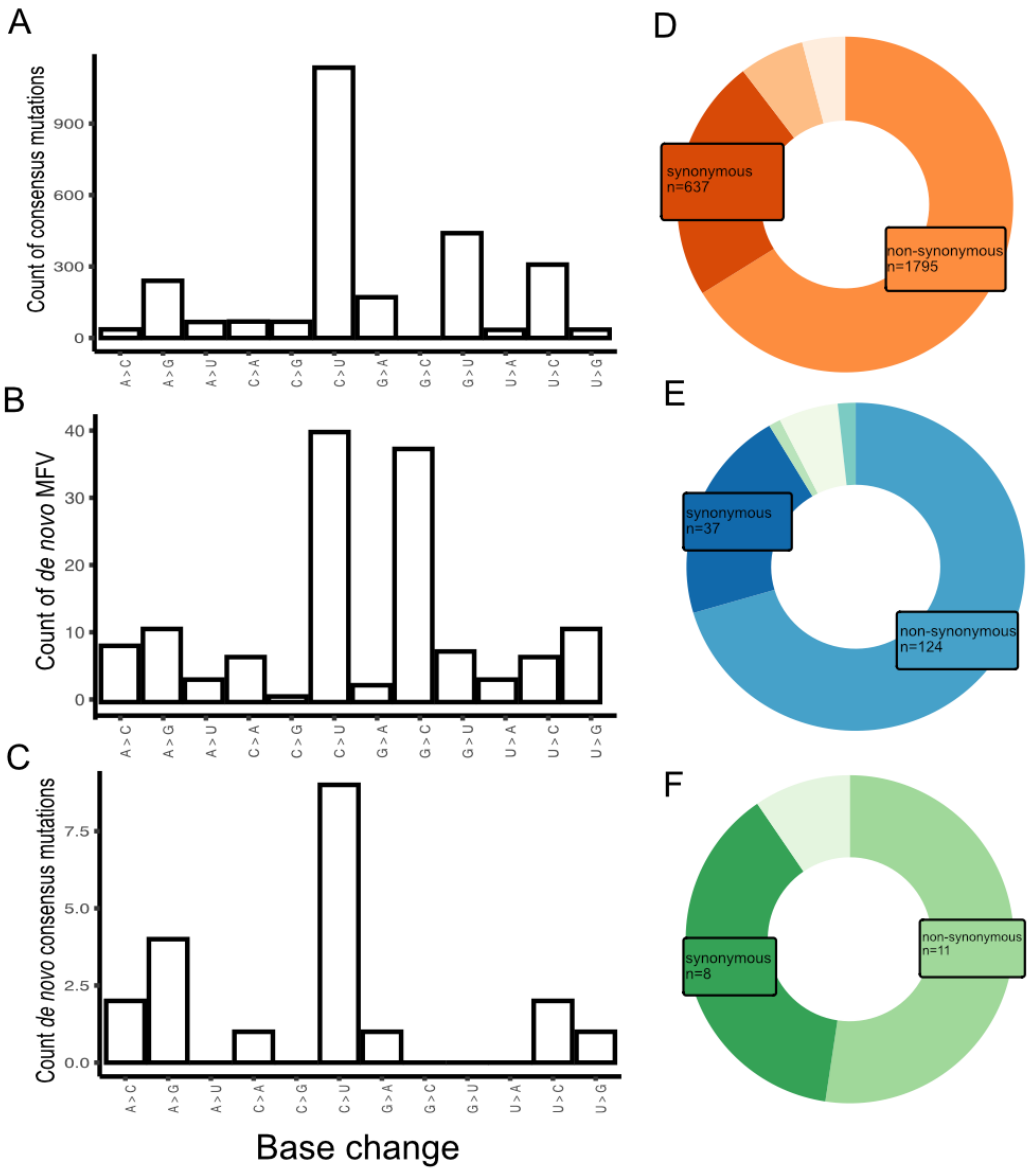


**Figure Description:** We quantified the mutational spectra during live virus neutralisation. (**A**) All consensus mutations detected against the reference genome (**B**). *De novo* MFV detected 72 hours post-neutralisation not contained in the inoculating virus (**C**) The *de novo* consensus mutations detected 72 hours post-neutralisation not detected as a fixed mutation or MFV in the inoculating virus. **(D)** Variant class of consensus mutations detected against the reference genome (indels n=112, noncoding n=170) **(E)** Variant class of *de novo* MFV detected 72 hours post-neutralisation (indels n=10, noncoding n=2, nonsense n=3) **(F)** Variant class of *de novo* consensus mutations detected 72 hours post-neutralisation (noncoding n=2)

**Key:** MFV – minority allele frequency variant

**Figure S3: Evolution of minority frequency variants in inoculating viruses 72 hours post-neutralisation.**

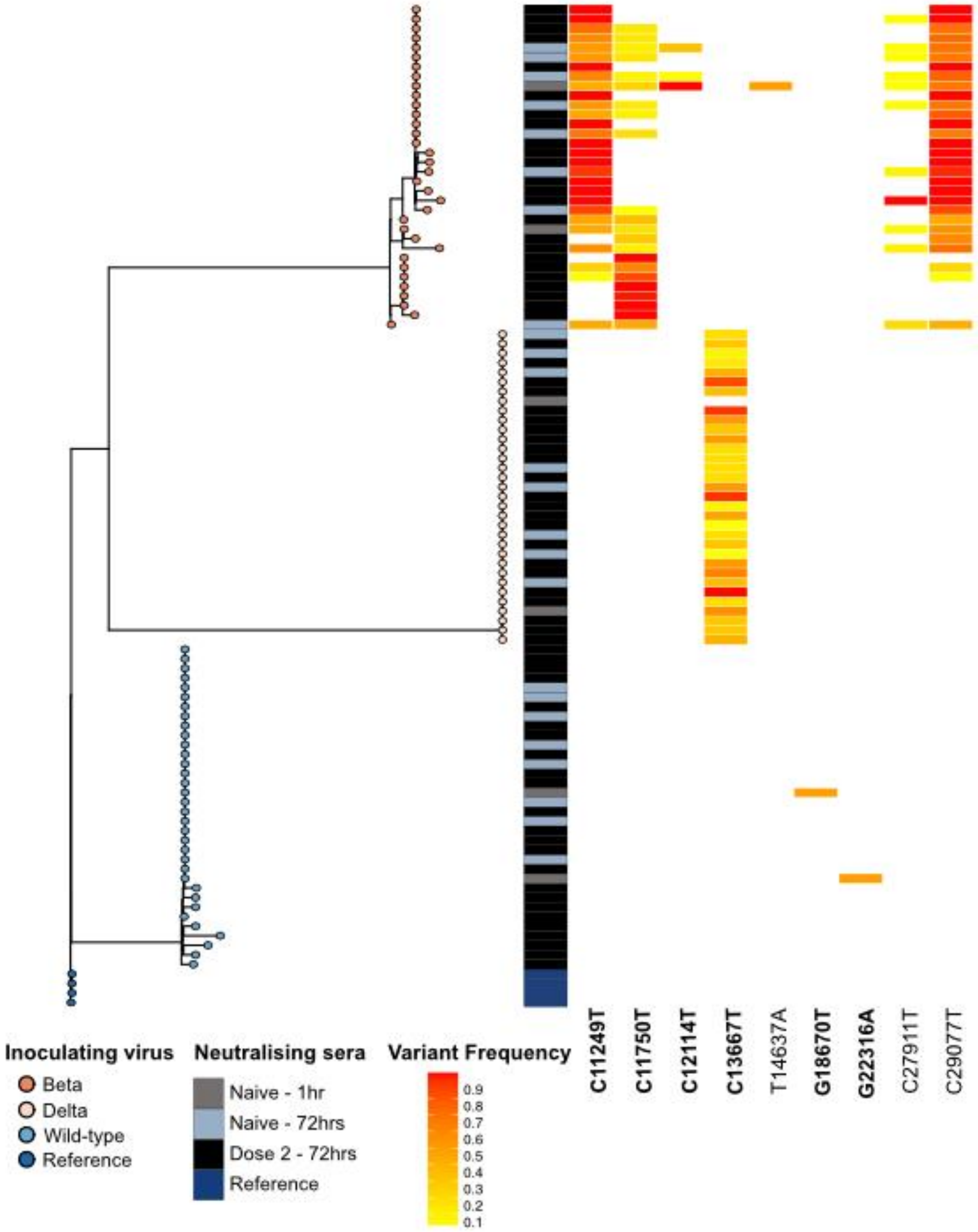


**Figure Description:** A maximum likelihood phylogeny of the 109 SARS-CoV-2 genomes generated in this study. Tree node colours indicate the lineage of the inoculating virus and the metabar highlight the time of sampling and the vaccine dose of the sera used. *De novo* minority allele frequency variants (MVF) which were detected in the sequence of each inoculating virus at specific genomic locations are shown in the heatmap. The read frequency of these sub-consensus variants is depicted by the colour scale, were a frequency of 0.1 is shown in yellow and a frequency of 0.9 is shown in red.

**Key:** Beta – (B.1.351) lineage; Delta - (B.1.617.2) lineage; Dose 2 – post the 2nd dose of Pfizer-BioNTech (BNT162b2) SARS-CoV-2 vaccination; Reference – reference SARS-CoV-2 genome (NCBI GenBank accession MN908947.3; Wild type – (A.2.2) lineage
